# Knowledge, attitudes and practices regarding rabies and its control among dog owners in Kigali city, Rwanda

**DOI:** 10.1101/500595

**Authors:** P. Ntampaka, P.N. Nyaga, J.K. Gathumbi, M. Tukei, F. Niragire

**Affiliations:** Department of Veterinary Medicine, University of Rwanda, Nyagatare, Rwanda; Department of Veterinary Pathology, Microbiology and Parasitology, University of Nairobi, Nairobi, Kenya; Department of Applied Statistics, University of Rwanda, Kigali, Rwanda

**Author notes:** Corresponding author (PN). These authors contributed equally to this work (P. Ntampaka, P.N. Nyaga). These authors also contributed equally to this work (J.K. Gathumbi, M. Tukei, F. Niragire).

## Abstract

**Background:** Rabies is a zoonotic viral disease that can occur in all warm-blooded mammals, including man [1]. Vaccinating dogs can protect people from contracting rabies [2]. Annual deaths due to rabies was estimated to 61000 worldwide [1], and Africa represented 35.2% of the deaths [3]. In Rwanda, rabies is a public health threat to the public [4], but the country does not have information on the disease [5].

**Methodology:** The present study aimed to assess knowledge, attitudes and practices (KAP) of rabies and its control among dog owners in Kigali city, Rwanda. We conducted a cross-sectional survey using a structured questionnaire among 137 dog owners randomly selected within each of the selected 9 study sites. A series of chi-square tests of association and binary logistic regression were used to determine the important factors associated with the response variables.

**Results:** The results showed that 99.5% of respondents could mention at least a host susceptible to rabies. Only 22.4 % and 21.3 % knew about canine and human rabies, respectively. Nearly 73.6% knew that human rabies can be transmitted through dog-bites and 99% could identify at least a clinical sign of canine rabies. Nealy 81.8% thought that regularly vaccinating dogs could prevent people from contracting dog-transmitted rabies. Only 43.1% and 26.3% were aware that clinical human and canine rabies are always deadly, respectively. Respondents who would observe a dog for some time, once it bites a man or an animal, represented 69%. Only 20.4% were familiar with cleaning dog-bites wounds with water and soap, before attending health care facilities. Few respondents owning dogs (20.6%) knew that puppies could receive rabies vaccination before the age of three months. Of respondents who owned vaccinated dogs, 78% were happy about the cost of rabies vaccination of dogs (Rwandan Francs 0-30,000). Nearly 57.9% had their dogs vaccinated at home by veterinarians.

Eighty-two (82%) percent of respondents received rabies information from neighbours, the media and public meetings. Logistic regression analyses indicated that none of respondents’sex, education level, and duration of dog ownership was statistically associated with their knowledge of rabies. The respondents who had kept dogs for 5-10 years were less likely to have as sufficient knowledge as those who had kept dogs for more than ten years (AOR=0.96). Male respondents were more likely to adopt a positive attitude (AOR=1.47) and have appropriate practice (A0R=1.40) towards rabies. The respondents who had completed at least primary education, were more likely to have appropriate practice of rabies (AOR=1.41).

**Conclusions:** This study identified gaps in the dog owners’ rabies knowledge of transmission, treatment and control. In addition, none of respondents’sex, educational level, and dog ownership length was statistically associated with their rabies knowledge. Overall, this study indicated that all categories of dog owners in Kigali city did not have good levels of rabies knowledge. Rabies interventions including awareness component in the studied population should be homogeneously improved.

## Introduction

Rabies is a zoonotic viral disease that can occur in all warm-blooded mammals, including man [1]. It is caused by a lyssavirus [6], *Lyssaviridae* contains 15 species, including rabies virus [7]. Rabies disease results in encephalomyelitis and is always deadly [2]. Worldwide, it was estimated that 61000 people die annually due to rabies [1] while post-exposure vaccination is given to over 15 million people [8]. In Africa, it was estimated that rabies kills 21476 people every year [3]. Annual livestock losses due to rabies was approximated to US$ 12.3 million [9–1]. Rabies virus is mainly transmitted through bites, but can also be transmitted through scratching and licking [10]. Transmission can also be through mucous membranes [11]. Human rabies is mainly caused by dog-transmitted rabies virus [12].

Worldwide, almost 100% of human rabies cases are transmitted by dogs [2]. Animal rabies is typically featured by sudden change in behaviour and progressive paralysis leading to death depending on the effect of rabies virus on brain [13). Clinical signs of canine rabies include biting without provocation, eating abnormal items, roaming, change in sound, hypersalivation [14). There are 6 stages that countries known to be endemic for rabies transmitted by dogs must go through to eliminate the disease. When rabies is likely to be present in a country and the country does not have information on the disease, that country is said to be at stage 0 [15]. Rwanda was reported to be at stage 0 [5). In Rwanda, majority of dogs owned by individuals are guard, although few are kept as pets or for business. Non-confined pet dogs outnumber confined ones and may interact with wildlife species and human dog-bites are predominantly done by the non-confined and stray dogs (Personal communication). According to [4], rabies is a serious public health threat to Rwanda. Between January and August 2016, 413 cases of human dog-bites were reported across Rwanda and one person died of rabies [16]. Controlling rabies in Rwanda is achieved through annual campaigns for vaccinating owned dogs and culling stray dogs [17].

According to Rwanda Agriculture Board, in 2016, dog population in Rwanda was estimated to 18117 including 11375 vaccinated against rabies and 2870 culled; the coverage of vaccination reached 62.7% [18]. Rwanda national veterinary laboratory is equipped to confirm suspicious cases of animal rabies, but poor collaboration between veterinary and medical personnel impacts on the surveillance and control as well as on awareness of the public on rabies [19]. Lacking information on real burden of rabies across mainland Africa, constitutes a challenge to its control and eradication [20]. Through upgrading public knowledge, KAP surveys help in changing attitudes and practices [21].

Recent published studies on awareness in most rabies endemic areas showed gaps in rabies knowledge. For example, a KAP survey in Haiti, indicated that 34% of medical professionals were familiar with using water and soap to clean bite wounds while 2.8% of them knew that rabies vaccination was part of post exposure prophylaxis [22]. A KAP survey in Ethiopia, showed that of interviewed pastoralists, 79.2% were unaware of how dogs got rabies while 23% of them did not know canine rabies symptoms [23].There are no published studies on rabies awareness among the population of Rwanda and it is not known whether Rwandese have knowledge of rabies. Thus, this study assessed dog owners’ knowledge, attitudes and practices of rabies disease and its control in Kigali city, Rwanda. It was hypothesized that dog ownership length did not impact knowledge, attitudes or practices of rabies in people owning dogs in Kigali city, Rwanda.

## Methods

### Study setting

The present study was carried out in Kigali, the capital city of Rwanda from September 2016 to February 2017. Rwanda is one of the landlocked countries located in Eastern Africa. Kigali City is one of the 5 provincial administrative entities, each subdivided into Districts, which in turn are subdivided into Sectors [24]. Kigali City is divided into three districts, namely Nyarugenge, Gasabo and Kicukiro.

**Fig 1.**
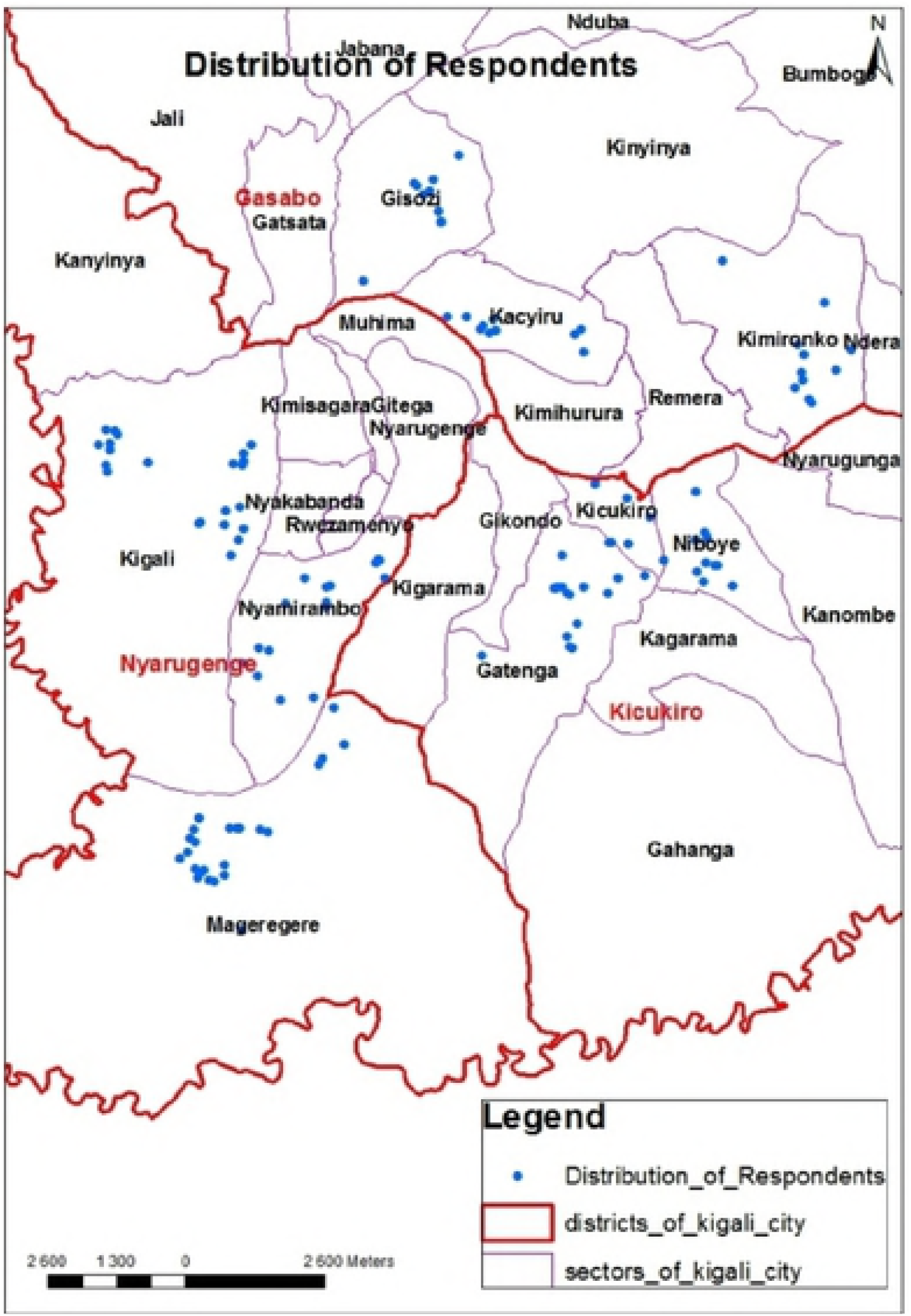
Distribution of the respondents across the nine study sectors. The figure illustrates the map for study setting and it was generated by authors. During data collection, a GPS was used to collect geographical information on each respondent household, and ArcGis10.2 software was used to produce the map from the GPS data. Names of three administrative districts of Kigali city are coulored red, while those of all administrative sectors of Kigali city are coulored black. Blue points symbolise the homes of respondents across the nine study sectors of which Mageragere, Nyamirambo and Kigali of Nyarugenge district, Gatenga, Niboye and Kicukiro of Kicukiro district, and Kacyiru, Kimironko and Gisozi of Gasabo district.

Kigali City was purposively selected on assumption that it harboured a large number of dogs, thus we expected to reach dog owners in a relatively limited time. The 2017 records in the three districts of Kigali city (Kicukiro, Nyarugenge and Gasabo) indicated that the dog population in Kigali city in 2016 was estimated to 2157 [25]. Considering that the dog population in Rwanda in 2016, was estimated to 18117 [18], the dog population in Kigali city in 2016, accounted for 12% of dog population in Rwanda. In the first stage, information from district-level registers on rabies vaccinations of dogs was used to select the nine 9 sectors across the 3 districts. In the second stage 137 respondents were systematically selected from dog owner’s population for each sector.

### Ethical standards

The Rwanda National Ethics Committee reviewed and approved this study (Review Approval Notice: No. 15/RNEC/2017), and dog owners signed certificates of consent prior to being interviewed. The certificate of consent is part of supporting document (“S1_File.pdf”). Distribution of the sample across selected administrative Sectors and Districts is shown (Table 1).

**Table 1.**
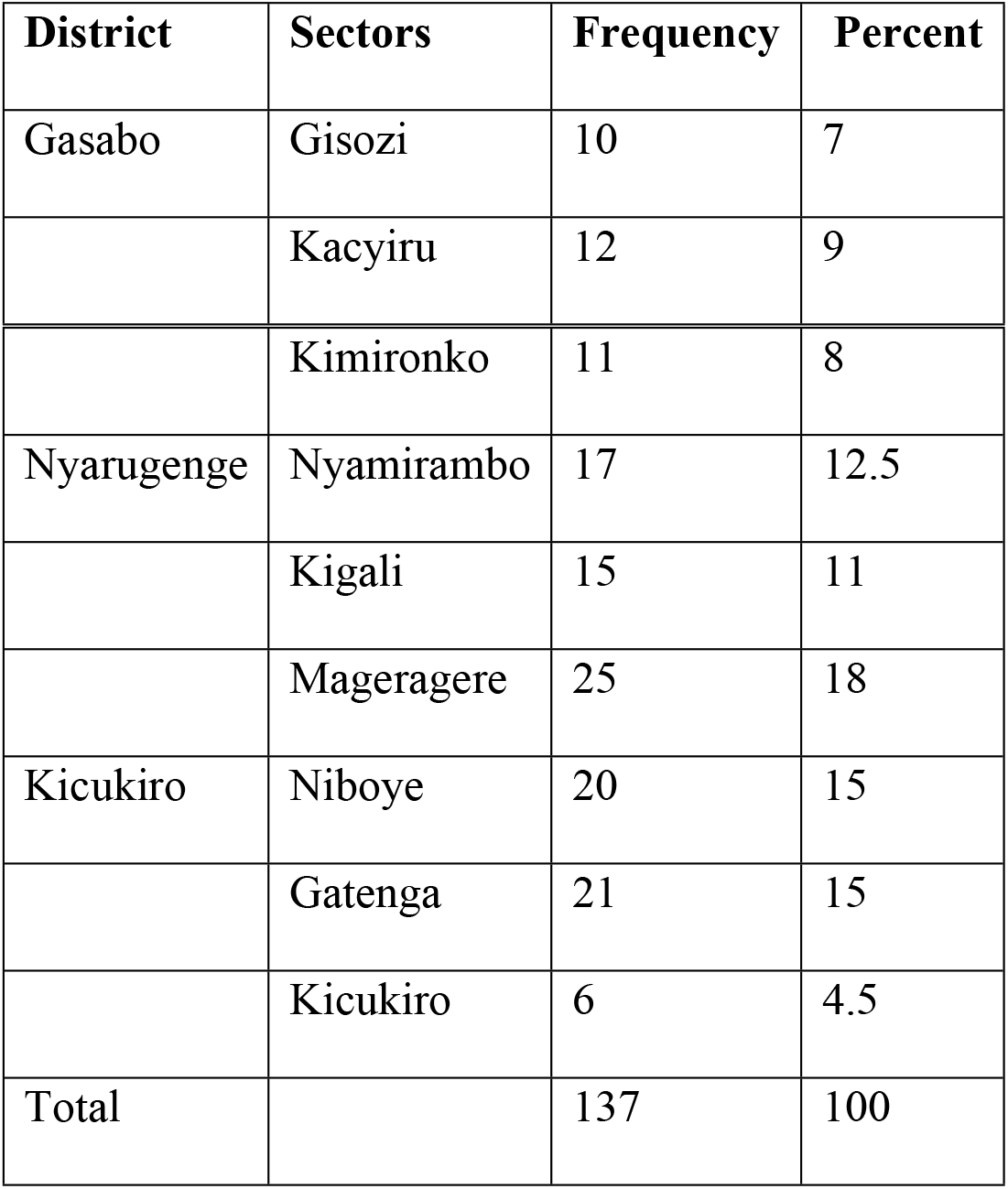
This is the Table 1 Distribution of the sample across selected Sectors and Districts.

| District | Sectors | Frequency | Percent |
| --- | --- | --- | --- |
| Gasabo | Gisozi | 10 | 7 |
|  | Kacyiru | 12 | 9 |
|  | Kimironko | 11 | 8 |
| Nyarugenge | Nyamirambo | 17 | 12.5 |
|  | Kigali | 15 | 11 |
|  | Mageragere | 25 | 18 |
| Kicukiro | Niboye | 20 | 15 |
|  | Gatenga | 21 | 15 |
|  | Kicukiro | 6 | 4.5 |
| Total |  | 137 | 100 |

Table 1 shows that respondents who lived in Nyarugenge district accounted for 41.5% while those from Kicukiro and Gasabo districts represented 34.5% and 24%, respectively.

### Selection and interview of respondents

Through collaboration with local authorities, dog owners were contacted by phone or in person. The questionnaire was pretested before administration and translated from English into Kinyarwanda (local language) during interview. The survey questionnaire is part of supporting document (“S2_File.pdf”).

### Analytical methods

During the interview, each respondent was given an identifying unique code that linked the respondents to their individual characteristics and responses during the data analysis process. Data organization and analysis were carried out using International Business Machines (IBM) Statistical Package for Social Sciences (SPSS) Statistics for Windows, version 20. In addition to frequency distributions analysis, tests of associations of knowledge, attitudes and practices regarding rabies with respondent’s individual characteristics (i.e., education level, sex, and length of dog ownership) were carried out using chi-square tests at 5% level of significance.

An index was constructed for each of the three components of interest, namely the level of knowledge, attitudes and practices towards rabies based on the respondents’ responses to corresponding questions using principal components factor analysis (PCFA) [26]. For data preparation for PCFA, items that were used to measure the three dimensions were transformed into indicator variables. First, the five items that were used to measure knowledge about rabies investigated whether the respondents knew: (i) *hosts who can suffer from rabies*; (ii) *how rabies can be transmitted between dogs and other animals*; (iii) *clinical signs of canine rabies*; (iv) *the prognosis for clinical canine rabies*; (v) *the most effective method for controlling canine rabies* (vi) *how best dog-mediated rabies can be controlled in humans*.

Respondent’s knowledge was classified as either *sufficient*, if they could provide correct answers to all the five items or *insufficient knowledge* otherwise. Second, respondent’s attitude towards rabies was measured using two items that aimed to find out: (i) *what they would wish to happen to the biting dog if it is caught*; and (ii) *what they can do before taking a colleague who is bitten by a dog to a health care facility*. The attitude was considered as either *positive*, if the respondents could indicate a correct attitude for the two items, or negative otherwise.

The respondents practices regarding rabies were assessed by investigating: (i) *the frequency (how often) respondent took their dog(s) to vaccination*; (ii) *how a veterinarian who previously vaccinated the respondent’s dog(s) carried and administered the vaccine*. Respondent’s practices were classified as either *appropriate*, if the respondent showed good practices vis-à-vis these items, or *inappropriate* or *bad* practices otherwise. All the indicators of respondent’s knowledge, attitudes or practices regarding rabies had two categories. Thus, binary logistic regression analyses were conducted for each indicator in order to identify and quantify net associations of each of the components with the respondents’ characteristics [27, 28–26]. A binary logistic regression analysis was conducted for each index of knowledge, attitude, and practice regarding rabies in order to determine the direction and extent of the effects of respondent’s individual characteristics, namely, sex, education level, length of dog ownership, and district of residence on the status of the three components (indices). All predictor variables were categorical and were all dummy coded.

## Results

### Demographic characteristics of the respondents

Overall, 137 respondents were interviewed including those owning dogs received rabies vaccination 107 (78.1%) or unvaccinated dogs 30 (21.9%). Of respondents, 65.7% were male while 34.3% were female. The respondents who did not get formal education represented 3.6% while those who finished primary and secondary school accounted for 30.7% and 28.5%, respectively. Approximately 37.2% finished or were receiving university education. Of interviewees, 43.8% and 22.6% had kept dogs for not more than 5 years and for 5-10 years, respectively while 33.6% had owned dogs for more than 10 years.

Of respondents, 57 (41.6%) lived in Nyarugenge district, while 47 (34.3%) and 33 (24.1%) dwelt in Kicukiro and Gasabo districts, respectively. Respondents obtained information on rabies from various sources, as shown in Table 2.

**Table 2.**
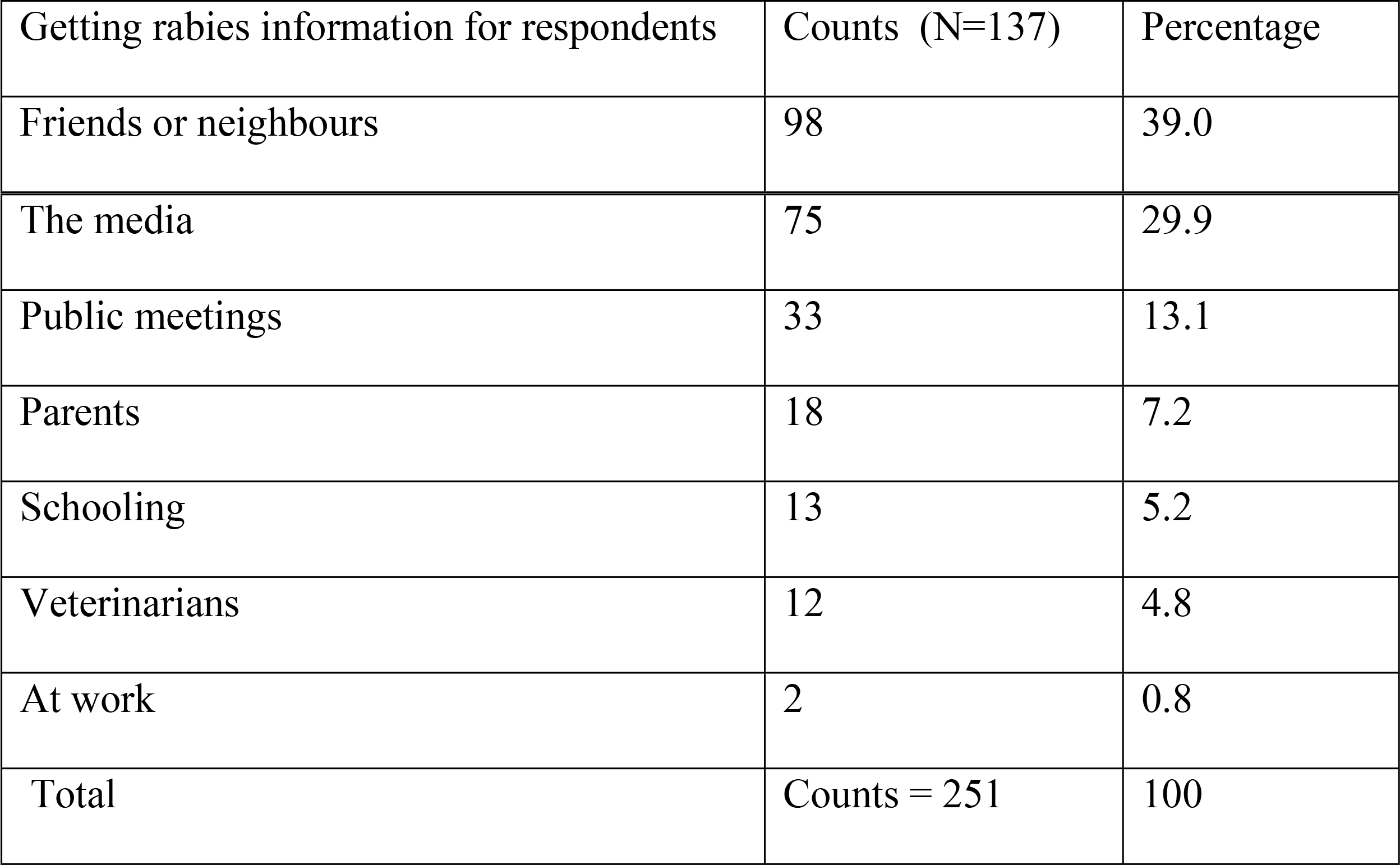
This is the Table 2 Sourcing rabies knowledge for respondents.

Table 2 shows that 82% of respondents sourced rabies information from neighbours, media and public meetings while 18% got the information from parents, school, veterinary personnel and workplace. The 0.8% who sourced rabies information from work included a medical pharmacist and a policeman, and they learnt about the disease through helping dog-bites patients or their relatives.

### Respondents’ knowledge of rabies

The awareness of respondents on the susceptible animal category is presented in Table 3.

**Table 3.**
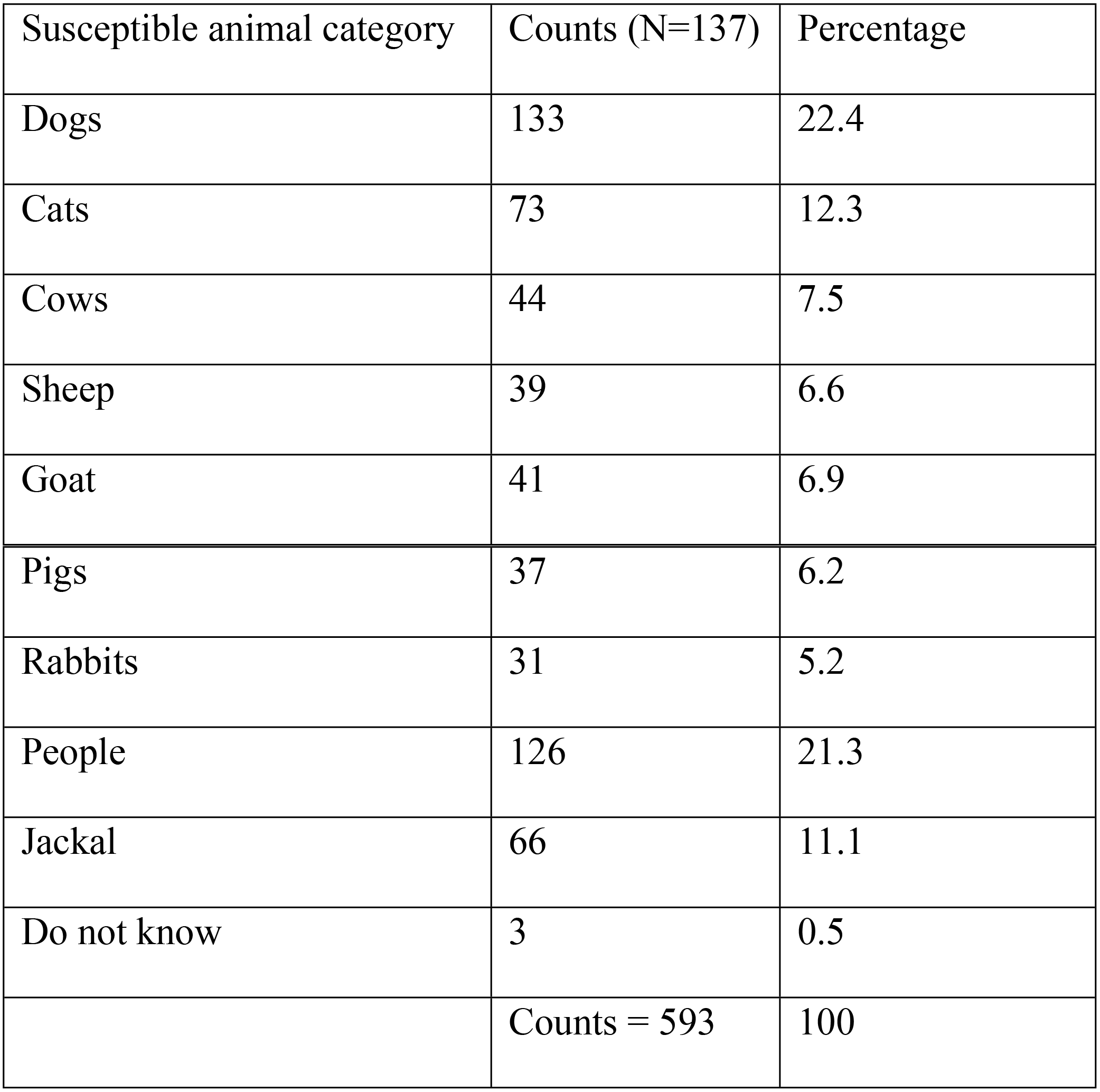
This is the Table 3 Participants knowledge of rabies susceptible hosts.

| Susceptible animal category | Counts (N=137) | Percentage |
| --- | --- | --- |
| Dogs | 133 | 22.4 |
| Cats | 73 | 12.3 |
| Cows | 44 | 7.5 |
| Sheep | 39 | 6.6 |
| Goat | 41 | 6.9 |
| Pigs | 37 | 6.2 |
| Rabbits | 31 | 5.2 |
| People | 126 | 21.3 |
| Jackal | 66 | 11.1 |
| Do not know | 3 | 0.5 |
|  | Counts = 593 | 100 |

Table 3 shows that 67.1% were aware of rabies in human, dogs, cats and jackals while 32.4% knew that farm animals (cows, sheep goats, pigs and rabbits) can have rabies. Approximately 0.5 did not have an idea on susceptibility of various hosts to rabies. Routes of rabies virus transmission to humans reported (Fig 2).

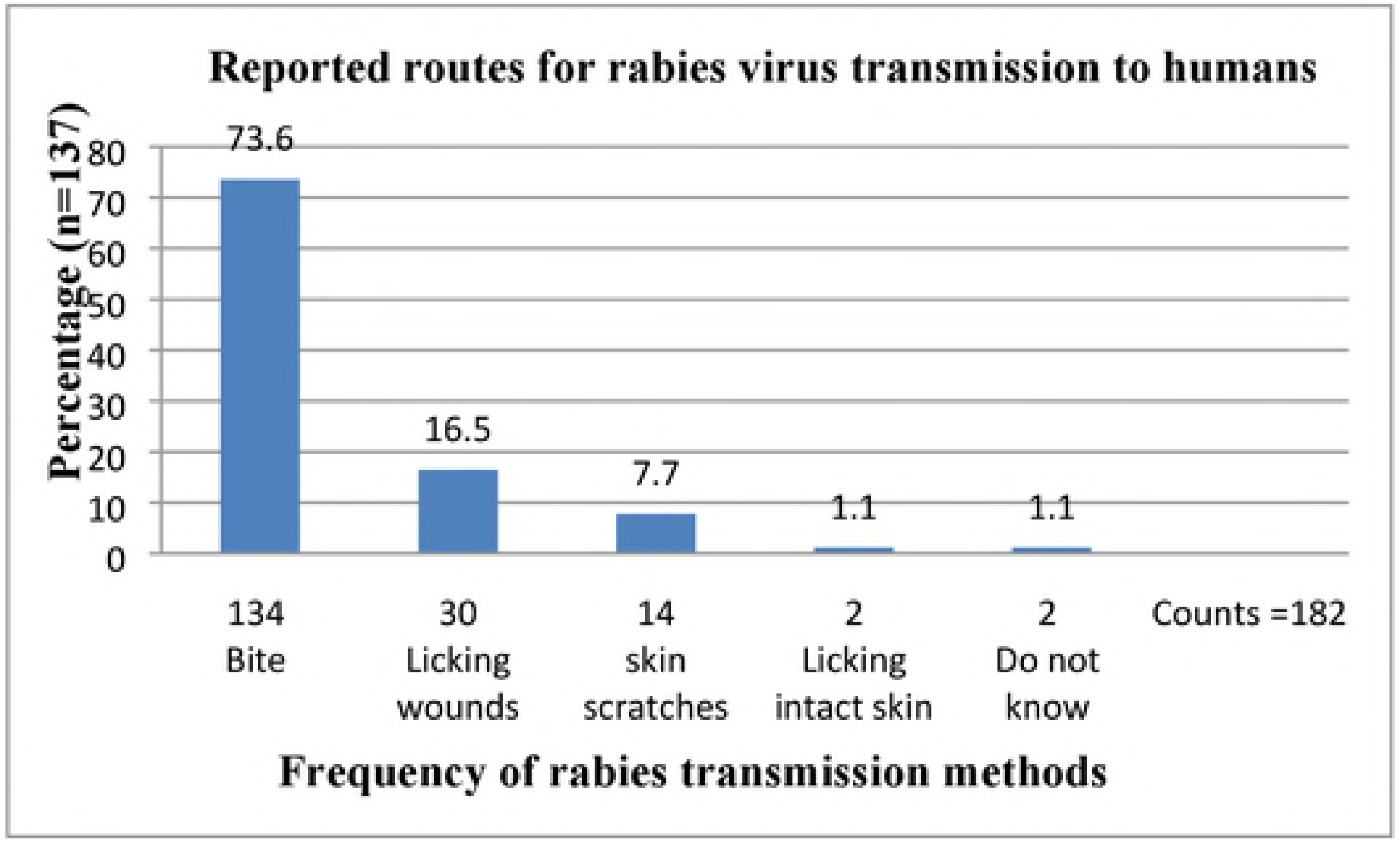
Reported routes for rabies virus transmission to humans. Fig. 2 shows that of 137 respondents, 73.6% knew that human rabies can be transmitted via dog-bites, while 16.5% and 7.7% were aware that humans can contract rabies through licking of wounds and skin scratches, respectively. Approximately 2.2% were unaware of transmission routes or believed in wrong route of transmission (licking intact skin) of how human rabies can be contracted. Respondents’ knowledge of animal rabies transmission is shown in Table 4.

Table 4 indicates that 85.4% of respondents knew how rabies can be transmitted between dogs and other animals (bites, licking of wounds and skin scratches), while 14.6% were misinformed (food, licking intact skin, coitus, inhalation) or unaware of transmission routes. Clinical picture of canine rabies as per respondents is shown in Table 5.

**Table 4.**
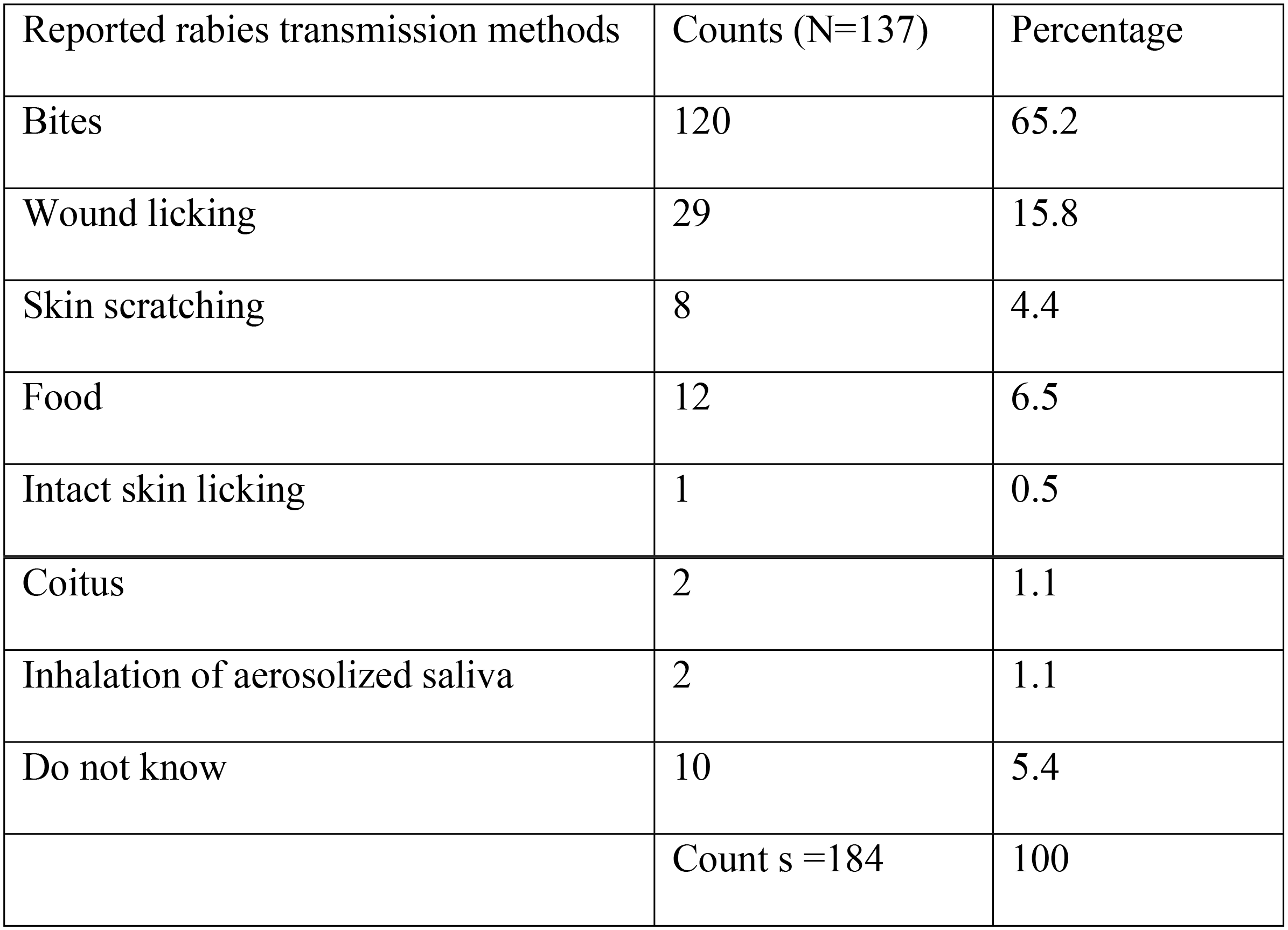
This is the Table 4 Respondents’ knowledge of animal rabies transmission.

**Table 5.** This is the Table 5 Manifestations of clinical rabies in dogs.

| Reported rabies clinical signs | Count (N=137) | Percentage |
| --- | --- | --- |
| Aggressiveness | 112 | 27 |
| Profuse salivation | 85 | 20 |
| Eating abnormal items | 43 | 10 |
| Difficulty in swallowing | 7 | 2 |
| Roaming over long distances | 97 | 23 |
| Change in sound | 31 | 7 |
| Dropping of the jaw | 41 | 10 |
| Do not know | 2 | 1 |
|  | Counts = 418 | 100 |

Table 5 shows that of respondents, 99% recognised at least a clinical sign of rabies in dogs while 1% was unaware of clinical signs of rabies. When asked about treating human rabies, 137 (42.3%) of respondents believed that human clinical rabies can be treated successfully, while 43.1% thought that it is always fatal, and 14.6% did not know whether it is treatable or fatal. Of the 137 study respondents, 54% believed that canine clinical rabies can be treated successfully, while 26.3% thought that it is always fatal, and 19.7% did not know whether it is treatable or fatal.

Of 137 respondents, 81.8% considered regular vaccination of dogs a method of choice for protecting people from getting rabies, while 18.2% believed that other methods would suffice to prevent and control human dog-mediated rabies. These methods included killing stray dogs (5.9%), educating the public (5.1%), completely restricting dogs (2.9%), immunising people at risk of developing rabies (2.9%), and treating people after exposure (1.4%).

### The respondents’ attitudes regarding motiveless human dog bites

Of 137 study respondents, 69% would keep a dog for some time once it bites a man or an animal to verify whether it was rabid or not, irrespective of the dog’s vaccination status. The percentage of respondents who would kill or release the dog immediately accounted for 15.5% for each category, regardless of dogs’ vaccination status and whether its owner was known. When asked about caring for a dog bite patient, 68.6% of respondents indicated they would take the patient to a hospital before they do anything.

Approximately 20.4% would clean the victim’s wounds with water or with both water and soap depending on availability; while 8% would cover the patient’s wound with dressings and bandages. Those who would put salt on the wound or clean it with 70% alcohol or povidone-iodine represented 1.5% for each.

### The respondents’ practices of rabies control

Only 20.6% of respondents who owned vaccinated dogs (n=107), vaccinated their puppies before they were 3 month old. The 79.4% who did not vaccinate theirs did not think dogs of such age could receive rabies vaccination. In practice, 57.9% and 41.2% of respondents had their dogs vaccinated by veterinarians at their homes and at site during campaign of vaccination, respectively.

Less than 1% had their dogs received rabies vaccination at a veterinary clinic. Of respondents (n=107), 86% and 12.1% indicated that, at time of vaccination, their veterinarians kept the vaccines with a cooler box and plastic bag (containing ice), respectively. Only 1.9% of respondents availed rabies vaccines home and then veterinarians administered them. This study showed that rabies vaccination fees varied between Rwandan Franc 0-30000; 78% of respondents were happy with the cost of vaccination while 22% were unhappy saying the cost was expensive. Respondents who owned unvaccinated dogs (n=30) indicated the main reasons for not vaccinating dogs included lacking information (39.6%), negligence (37.2%) and inadequate knowledge of rabies (11.6%). Vaccination fees and vaccination sites located far from home were the other reasons for having unvaccinated dogs, representing 9.3% and 2.3%, respectively.

### Results of multivariable analyses

For each of the indexes of the status of knowledge, attitudes and practices regarding rabies, a simple binary logistic regression model was fitted to data for each of the selected individual dog owner’s characteristics. The adjusted odds ratios (AOR) from the series of the three logistic regression analyses are shown in Table 6.

**Table 6.** This is the Table 5 Adjusted Odds Ratios (AOR) with 95% Confidence Intervals (CI)

| Characteristics | Category | Knowledge | Attitude | Practice |
| --- | --- | --- | --- | --- |
| District | Gasabo | *Reference* | *Reference* | *Reference* |
|  | *Kicukiro* | 0.59(0.23,1.54) | 0.65(0.18,2.33) | 0.55(0.21, 1.44) |
|  | *Nyarugenge* | 0.42(0.15,1.18) | 1.74(0.46,6.51) | 0.98(0.32, 2.98) |
| Sex | *Female* | *Reference* | *Reference* | *Reference* |
|  | *Male* | 0.60(0.27,1.34) | 1.47(0.49,4.42) | 1.40(0.62, 3.13) |
| Education level | *Tertiary* | *Reference* | *Reference* | *Reference* |
|  | *No education* | 2.12(0.27,16.45) | 0.41(0.03,5.40) | 0.71 (0.09, 5.66) |
|  | *Primary* | 0.97 (0.37,2.54) | 0.50(0.14,1.85) | 1.42 (0.52,3.88) |
|  | *Secondary* | 1.24 (0.51,3.05) | 0.59(0.18,1.97) | 1.51(0.59,3.86) |
| Length of dog ownership | *> 10 years* | *Reference* | *Reference* | *Reference* |
|  | *< 5 years* | 1.23 (0.54, 2.79) | 0.39(.14, 1.11) | 0.97(0.42,2.27) |
|  | *5-10 years* | 0.96 (0.37, 2.50) | 0.35(.10, 1.26) | 1.46(0.52,4.13) |
| Constant |  | 2.33 | 0.38 | 1.53 |

Table 5 reveals that none of the predictor variables, namely respondent’s sex, education level; district of residence or the length of dog ownership was statistically associated with the respondent’s knowledge about rabies. Similarly, there was no statistically significant association of the status of dog owner’s attitudes or practices regarding rabies with any of the selected predictor variables. However, there were different relationships between the status of the respondent’s knowledge, attitude and practice and the selected classes of dog owner’s. First, the knowledge about rabies was more likely to be sufficient among the less educated dog owners compared to those who completed or were still pursuing tertiary education. The odds of sufficient knowledge were more than twice among those without formal education, and 24% higher among those with secondary education compared to those with tertiary education. The odds of sufficient knowledge were 40% lower among male compared to female dog owners. Those who had kept dogs for 5-10 years were less likely to have as sufficient knowledge as those who had kept dogs for more than ten years (AOR=0.96). However, those who had owned dogs for at most five years (<5years) were more likely to have sufficient knowledge about rabies (AOR=1.23).

The odds of sufficient knowledge among dog owners in districts of Kicukiro and Nyarugenge were respectively 41% and 58% lower compared to those living in Gasabo District. Second, the respondent’s attitude towards rabies was more likely to be positive in male respondents (AOR=1.47). The odds of positive attitude were more than 40% lower among those who completed any level of education other than tertiary education. Similarly, those who had owned dogs for more than 10 years were more likely to show positive attitudes compared to other dog owners. In particular, the odds of positive attitude towards rabies were 65% lower for those who had kept dogs for 5-10 years compared to those who had owned dogs for more than 10 years. Third, the respondent’s practices regarding rabies were more likely to be appropriate for male dog owners (AOR=1.40), living in Gasabo, who had completed at least primary education (AOR=1.41), or who had kept dogs for at least five years (AOR=1.46).

## Discussion

The present study aimed to investigate dog owners’ knowledge, attitudes and practices of rabies disease and its control in Kigali City, Rwanda. The findings showed that 43.7% of respondents were aware of canine and human rabies while 2.2% did not know how humans can contract rabies. Approximately 14.6% were unaware of how rabies can be transmitted between dogs and other animals. Around 18.2% of respondents thought that killing stray dogs, educating the public, completely restricting dogs, vaccinating people at risk of having rabies, and treating people after exposure would suit controlling and preventing dog-mediated human rabies. Nearly 43.1% and 26.3% of respondents knew that once clinical signs of rabies appear, the disease is always fatal in humans and dogs, respectively. Of respondents, 20.4% knew that dog-bites wounds should be washed with soap and clean water before attending a hospital while 31% did not think that, a biting dog might be confined for some time to assure it was not rabid.

Only 20.6% of the respondents knew that puppies can receive rabies vaccination before the age of 3 months old. It is important to fill in these gaps in rabies knowledge because, for example, knowing how dogs can potentially contract rabies would help understand that timely vaccination of dogs is the method of choice for controlling and preventing canine rabies. Being aware that rabies is incurable would show that the only solution is vaccinating dogs. Knowing what should be done after a dog bites a person or an animal would help understand that the dog should be quarantined for some time to assure it is not rabid and that dog-bites wounds should be cleaned before attending a hospital (changing from negative towards positive attitudes).

We found that only 43.7% of the respondents were aware of human and canine rabies, and this was lower than 70% of the respondents reported by [29]. Our findings showed that 73.6% of the respondents knew that human rabies can be transmitted through dog-bites. This was comparable to 73% of the respondents reported by [22] who knew that human rabies can be transmitted through biting by a rabid animal. Our findings showed that 85.4% of the respondents were aware of modes through which rabies can be transmitted between dogs and other animals. The percentage exceeded 72.6% of interviewees reported by [30] who were aware that rabies can be contracted through bites, scratches and exposing open wounds to saliva. The 14.6% of our respondents who were misinformed or unware of how animal rabies can be transmitted were lower than 84% who believed that exposing infected saliva to wounds or intact skin can result in rabies and 32% who thought that the disease can be transmitted through inhalation as it was reported by [31].

In this study, 99% could identify at least a clinical sign of canine rabies and aggressiveness, roaming aimlessly and salivation were the most known signs. A study conducted by [32] in Philippines found that 69% of the households were familiar with at least a symptom of canine rabies and that the three famous symptoms were salivation, weakness and change in behaviours. The 81.8% of our respondents who indicated that regularly vaccinating dogs would be the method of choice for protecting people from getting dog-mediated rabies was higher than 41.7% of the interviewees who knew that vaccination of dogs can help prevent them from getting rabies as it was reported in Ethiopia [33]. Regarding knowledge of rabies among the respondents, only 43% and 26% knew that clinical human and canine rabies are almost always fatal, respectively. A previous study [29] found that 63% of the respondents who were aware that, once rabies manifests clinically, it is always deadly.

According to [34], rabies cannot successfully be treated, once it manifests clinically. This study showed that respondents without formal or with secondary education were more likely to have good knowledge of rabies than those who finished primary or tertiary education. Our findings disagreed with those of [35,36–37) who found that respondents’ knowledge of rabies increased steadily from non-formal to tertiary education. The fact that all the respondents owned dogs and that the prime source of rabies information was neighbours could be an indication that dog owners interacted to share knowledge of rabies regardless of their education level. We found that the odds of sufficient knowledge were 40 % lower among male compared to female dog owners. A study conducted by [37] found that sex did not influence knowledge of rabies among respondents. We found that the length of dog ownership was associated with knowledge of rabies among dog owners. A study conducted by [38] in Kenya revealed that owning a dog for a long time for the respondents could make them realise the benefits of vaccinating their dogs regularly. This study revealed that the three leading sources of rabies information for the respondents were neighbours, radio and television as well as public meetings accounting for 76.8%. A study carried out by [23] comprising urban and pastoralist respondents in Ethiopia revealed that 92.1% of the urban interviewees received rabies information from families whereas a study by [39] found that 86.6% of the respondents sourced rabies information from neighbours. We found only 20.4% of respondents would use soap or both soap and water depending on availability to clean wounds caused by dog-bites. The 20.4% was lower than 43.07% of victims bitten by dogs who used soap and water to clean wounds before going to hospitals in India [40], it however exceeded 5% of interviewees who knew that dog-bite wounds should be cleaned before seeking medical care reported by [29].

According to [14], the emergency treatment of choice against rabies virus consists of washing and flushing human animal-bite wounds immediately with soapy water for around a quarter hour. We found that only 20.6% of the respondents vaccinated their puppies against rabies before the age of 3 months. According to [41], Majority of canine population across Africa consists of puppies younger than 3 months. Our results showed that the respondents who had owned dogs for more than 10 years were more likely to exhibit positive attitude towards rabies than those who had kept dogs for fewer years. Respondents’ practices of rabies were more likely to be influenced by sex, education level, and length of dog ownership.

## Conclusion

This study identified gaps in the dog owners’ rabies knowledge of transmission, treatment and control. In addition, none of respondents’sex, educational level, and dog ownership length was statistically associated with their rabies knowledge. Overall, this study indicated that all categories of dog owners in Kigali city did not have good levels of rabies knowledge.

Rabies interventions including awareness component in the studied population should be homogeneously improved.

## Acknowledgements

The authors would like to thank the leaders of Kicukiro, Gasabo and Nyarugenge districts in Kigali City for allowing data collection. Sincere thanks to the dog owners for frank collaboration during the interviews.

## Supporting information

**S1 Fig. This is the S1 Fig Title. Distribution of the respondents across the nine study sectors**

**S2 Fig. This is the S2 Fig Title. Reported routes for rabies virus transmission to humans**

**S1 File. This is the S1 File Title. Certificate of consent for respondents**

**S2 File. This is the S2 File Title. The survey questionnaire**

